# Partial ablation of the orexin field induces a sub narcoleptic phenotype in a conditional mouse model of orexin neurodegeneration

**DOI:** 10.1101/234765

**Authors:** Sarah Wurts Black, Jessica D. Sun, Pamela Santiago, Alex Laihsu, Nikki Kimura, Akihiro Yamanaka, Ross Bersot, Paul S. Humphries

**Author notes:** Communicating author: Sarah Wurts Black, Ph.D., Reset Therapeutics 260 Littlefield Ave, Suite 200 South San Francisco, CA 94080, USA.

## Abstract

Narcolepsy type 1 (Na-1) and 2 (Na-2) are characterized by an inability to sustain wakefulness and are likely caused by degeneration of orexin neurons. Near complete orexin neurodegeneration depletes orexin-A from the cerebrospinal fluid and produces Na-1. The pathophysiology of Na-2 is less understood, but has been hypothesized to be due to less extensive loss of orexin neurotransmission. The *orexin-tTA; TetO diphtheria toxin A* mouse allows conditional control over the extent and timing of orexin neurodegeneration. To evaluate partial ablation of the orexin field as a model of Na-2, orexin-A positive cell counts and sleep/wake phenotypes (determined by piezoelectric monitoring) were correlated within individual mice after different protocols of diet-controlled neurodegeneration. Partial ablations that began during the first 8 days of study were 14% larger than partial ablations induced during the last 8 days of study, six weeks later and prior to sacrifice of all mice, suggesting orexin-A positive cell death continued despite the resumption of conditions intended to keep orexin neurons intact. Sleep/wake of mice with 71.0% orexin-A positive cell loss, initiated at the beginning of study, resembled that of orexin-intact controls more than mice with near complete neurodegeneration. Conversely, mice with 56.6% orexin-A positive cell loss, created at the end of study, had sleep/wake phenotypes that were similar to those of mice with near complete orexin-A positive cell loss. Collectively, these results suggest that compensatory wake-promotion develops in mice that have some critical portion of their orexinergic system remaining after partial ablation.

**Statement of significance:** The pathophysiology of narcolepsy type 2 is poorly understood but has been hypothesized to be due, at least in part, to degeneration of a smaller proportion of the orexin neuronal field than occurs in narcolepsy type 1. To evaluate a transgenic mouse model of narcolepsy type 2, we correlated the sleep/wake phenotypes of individual, male and female adult mice that received diet-induced conditional ablations of orexin neurons with their orexin cell counts. Using a translatable measure of narcolepsy sleepiness severity, we demonstrated that compensatory wake-promoting responses developed in mice concurrent with progressive orexin neurodegeneration. These results provide important details necessary for preclinical drug discovery for therapeutic areas characterized by orexin insufficiency, such as narcolepsy, Parkinson’s disease, and other neurodegenerative disorders.

## Introduction

The sleep disorder narcolepsy features excessive daytime sleepiness (EDS), rapid-eye-movement (REM) sleep abnormalities, disturbed nighttime sleep, and is caused by diminished orexin (also known as hypocretin) neurotransmission.^1^ Insufficient orexin-A levels in the cerebrospinal fluid (CSF) are thought to be due to autoimmune-related destruction of the sole population of neurons that produce orexin.^2–4^ Patients with narcolepsy type 1 (Na-1) have low CSF orexin-A levels, cataplexy (a sudden, reversible loss of muscle tone triggered by emotions), or both. Narcolepsy type 2 (Na-2) is based on exclusion of Na-1 criteria, and its etiology is less understood than the pathology underlying narcolepsy with cataplexy. However, a limited postmortem study suggests Na-2 can be caused by 33% orexin cell loss vs. the ~90% loss observed in patients with Na-1.^5^ Levels of CSF orexin-A that are between normal and the Na-1 cutoff value have been associated with Na-2 in a subset of patients.^6,7^ Both types of narcolepsy are diagnosed, in part, by EDS that is determined by the inability of patients to sustain long periods of wakefulness during the Maintenance of Wakefulness Test (MWT) and to fall asleep with short latency on the Multiple Sleep Latency Test (MSLT).

Two transgenic mouse models have been created that recapitulate the orexin neurodegeneration and arousal state boundary dysregulation observed in Na-1. The *orexin/ataxin-3* (Atax) mouse exhibits constitutive, postnatal loss of 80-99% of orexin neurons^8–11^ and express a range of narcolepsy symptom severity.^1^ The *orexin-tTA; TetO diphtheria toxin A* (DTA) mouse uses the double transgenic Tet Off (TetO) system to allow conditional ablation of orexin neurons.^12^ Administration of doxycycline (DOX) to these mice blocks the tetracycline transactivator from binding to the TetO binding site and disables the expression of the DTA neurotoxin. Thus, orexin neurodegeneration can be induced by removal of doxycycline (DOX(−)). Withdrawal of dietary DOX (100 μg/g chow) for 1.5, 3.5, 7, 14 and 28 days followed by resumption of DOX has been shown to induce 36%, 64%, 86%, 95%, and 97% orexin cell loss, respectively.^12^ Longitudinal study by the same research group on a different set of DTA mice on continual DOX(−) has shown fragmentation of wakefulness during the dark period at 1 week DOX(−) that progressively worsened over the next 3 weeks. During this time, cataplexy began at 2 weeks DOX(−) and increased in frequency over the next 5 weeks or more. Thus, conditional control of orexin neurodegeneration in DTA mice has been proposed to model either Na-1 or Na-2, depending on the extent of the ablation. However, to validate partial ablation of the orexin field as a model of Na-2, the precision of the DOX(−) timing to reliably induce a certain percentage of orexin cell loss that corresponds to a translatable phenotype of Na-2, all within the same animal, must be determined.

Recently, a translational test of the ability to sustain long bouts of wakefulness has been developed for mice using piezoelectric monitoring of sleep/wake.^13^ The PiezoSleep narcolepsy screen measures the ratio of time spent in long (≥ 32 min) wake bouts to time spent in short (< 8 min) sleep bouts to yield a “wake maintenance score.” Sleep/wake is sampled during the first half of the dark period, when sustained wakefulness predominates in wild type mice, to allow narcolepsy symptom severity to be assessed.

Here, orexin cell counts and sleep/wake phenotypes were correlated within individual DTA mice after partial ablation of the orexin neuronal field and sex differences were determined. The effectiveness of the Tet Off system to maintain intact orexin neurons after partial ablations induced by DOX(−) was evaluated. The stability of the sleep/wake phenotypes over time that correspond to different orexin neurodegeneration protocols were also assessed. These results further define the parameters that are essential to the validate the DTA model beyond Na-1.

## Materials and Methods

All experimental procedures were approved by the Institutional Animal Care and Use Committee at Reset Therapeutics and were conducted in accordance with the principles set forth in the *Guide for Care and Use of Laboratory Animals*.

### Animals

Male and female orexin-tTA mice were bred with B6.Cg-Tg(tetO-DTA)1Gfi/J mice (both strains from Nagoya University, Nagoya, JP) to produce *orexin-tTA; TetO diphtheria toxin A* (DTA) mice that were hemizygous for both transgenes.^12^ Male and female C57BL/6NTac were used as wild type (WT) controls in Study 1. All DTA and WT mice were bred at Taconic Biosciences, Hudson, NY. All breeders and DTA mice were given doxycycline (DOX) chow (200 μg/g chow, TD.98186, Envigo, Madison, WI), to deliver a daily dose of approximately 0.6 mg DOX, until they were switched to standard chow at the start of experimental orexin neurodegeneration. Male and female hemizygous transgenic B6.Cg-Tg(HCRT-MJD)1Stak/J mice (Atax) and wild type (WT) littermates (JAX stock #023418, Jackson Labs, Bar Harbor, ME) were used in Study 2 as comparator groups that represent a standard model of narcolepsy^8^ and controls. In Study 2, animals were individually housed at 22-25 °C, 30-70% relative humidity, and LD12:12 with *ad libitum* access to food, water and nesting material.

### Study Protocols

In Study 1, the effects of shipping conditions on DOX consumption and orexin neurodegeneration were assessed. Sixty DTA mice and 8 WT mice were air shipped in single-sex, group-housed compartments (*N* = 4-5/compartment) from Taconic in Hudson, NY to South San Francisco, CA. The WT mice (4 males and 4 females) were shipped with standard chow. Of the DTA mice, 15 males and 15 females were shipped with DOX chow; the remaining 15 males and 15 females were switched to standard chow at the time of boxing for shipment. DTA mice were weighed immediately prior to packing and upon arrival. The percentage of weight gained or lost during transit was used as a proxy measure of food consumption. The 8 WT mice, 8 DTA mice shipped with DOX chow, and 8 DTA mice that had been switched to standard chow were sacrificed for immunohistochemistry 48-50 h after they were packed for shipment (ZT4-6). Half of the DTA mice in each DOX group used for histology were female. All mice used for histology remained in their original shipping compartments until the time of sacrifice. The remainder of the DTA mice were retained for other experiments, including Study 2.

In study 2 (Figure 1), the relationship between the size and timing of partial ablation of the orexin neuronal field and sleep/wake phenotype was determined. Twenty-five DTA mice were randomly assigned into groups based on DOX diet schedule; males and females were represented across groups and individually housed upon arrival for the duration of study. Mice were either: (1) maintained on DOX chow for 7 weeks (DOX(+)7wk) to keep orexin neurons intact (*N* = 4), (2) switched to standard chow for 8 days and then resumed DOX(+) for 6 wk (DOX(−)8d (RD)6wk) to induce partial orexin neurodegeneration during the first study week (*N* = 6), (3) switched to standard chow 8 days prior to sacrifice (DOX(+)6wk DOX(−)8d) to induce partial orexin neurodegeneration during the last study week (*N* = 7), or (4) maintained on standard diet without doxycycline for 7 weeks (DOX(−)7wk) to induce full orexin neurodegeneration (*N* = 8). Sleep/wake was assessed by piezoelectric monitoring during Study Weeks 4 and 7. Mice were euthanized for immunohistochemistry at the end of the Week 7 recording (ZT6-8). Previously collected piezoelectric recordings from Atax mice (*N* = 31) and their WT littermates (*N* = 17), aged 15.4 ± 0.3 weeks and 29.4 ± 0.3 g, served as a reference as the standard model comparator groups, as described.^13^

**Figure 1.**
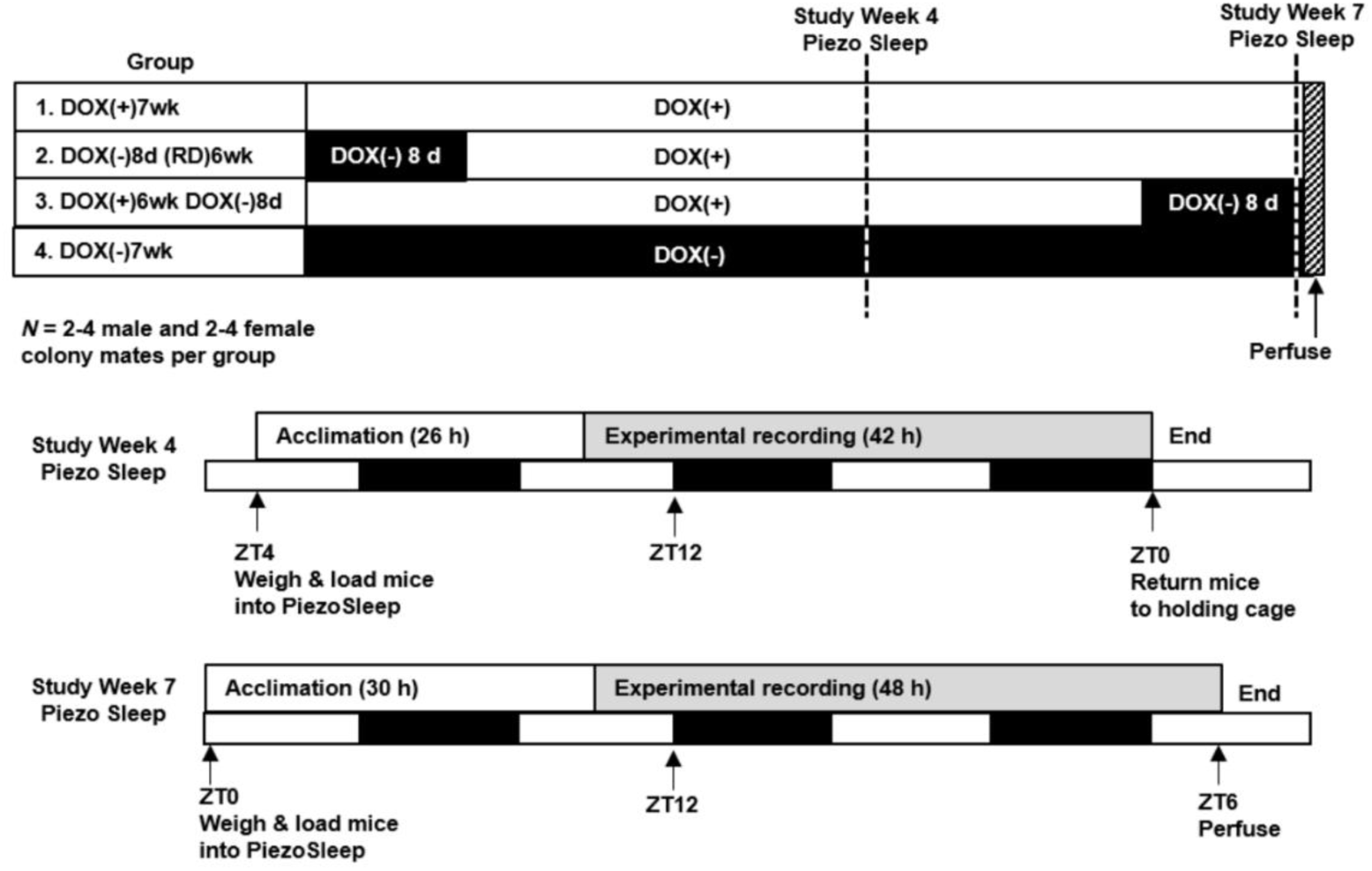
Study 2 protocol. *Orexin/tTA; TetO diphtheria toxin A* (DTA) colony mates were randomly assigned into study groups based on targeted orexin neuronal field ablation size and timing; males and females were represented across groups. DTA mice were either (1) maintained on doxycycline for 7 weeks (DOX(+)7wk) to keep orexin neurons intact (*N* = 2 males, 2 females), (2) subjected to removal of doxycycline for 8 days and resumption of DOX(+) for 6 wk (DOX(−)8d (RD)6wk) to induce partial orexin neurodegeneration during the first study week (*N* = 4 males, 2 females), (3) subjected to removal of doxycycline 8 days prior to sacrifice (DOX(+)6wk DOX(−)8d) to induce partial orexin neurodegeneration during the last study week (*N* = 4 males, 3 females), or (4) maintained without doxycycline for 7 weeks (DOX(−)7wk) to induce full orexin neurodegeneration (*N* = 4 males, 4 females). Sleep/wake was measured by piezoelectric monitoring at study weeks 4 and 7. For each sleep/wake recording, mice were moved from home cage to PiezoSleep cage for 20-24 h acclimation followed by experimental sampling for 30-48 h. At the end of the recording on study week 7, mice were sacrificed for histology.

### Immunohistochemistry

In Study 1, brains from DTA mice (8 males and 8 females) and wild type mice (4 males and 4 females), all aged 8.3 ± 0 weeks (22.0 ± 0.7 g) were used. In Study 2, brains from DTA mice (14 males and 11 females) all aged 15.1 ± 0.1 weeks (26.1 ± 0.9 g) were used. Mice were sacrificed with 0.1 mL Beuthanasia-D (Henry Schein, Dublin, OH) and transcardially perfused with 0.4 % paraformaldehyde. Brains were harvested, post fixed overnight, and shipped in phosphate buffered saline to NeuroScience Associates (Knoxville, TN), where all histology and cell counts on tissue from DTA and WT mice were performed. The brains were embedded together per study in a single gelatin matrix block. The multi-brain blocks were freeze-sectioned at 35 μm with brains in the coronal plane from approximately -1.23 to - 1.91 relative to Bregma. Every 4^th^ section was retained for the detection of orexin-A positive neurons. After blocking in normal donkey serum (5% in 0.3% triton in PBS), sections were incubated overnight at room temperature with goat anti-donkey orexin-A antibody (SC-8070, Santa Cruz Biotechnology, CA) at 1:2000 dilution. After washing with phosphate-buffered saline, sections were incubated for 2 hours at room temperature with biotinylated donkey anti-goat (Jackson Immunoresearch, West Grove, PA) at 1:500 dilution in blocking solution. After further washes, sections were developed with the avidin-biotin-peroxidase system using diaminobenzidine as the chromagen (Vector Laboratories, Burlingame, CA). Sections were mounted onto slides, and anatomically matched sections between brains that represented the anterior, mid, and posterior regions of the orexin field, approximately -1.4, -1.5, and -1.6 mm from Bregma, respectively, were identified for cell counting. A 1.4 mm^2^ counting box was delimited 0.3 mm from midline and 0.2 mm dorsal to the base of the 3^rd^ ventricle. Counts were performed bilaterally on these sections using Fiji Image J automated for thresholding and particle analysis of positively stained objects using a watershed filter and a measure of circularity on a 0 to 1 scale. Comparison of manual counts vs. automated counts on brains from WT and DTA mice with either no or partial ablation indicated 96-99% agreement. Disagreements mostly resulted from a few cells missed by automated counts because of overlapping somas. Immunohistochemistry for orexin-A on free-floating brain sections and manual cell counting methods for Atax mice (*N* = 13) and WT littermates (*N* = 14), aged 16.9 ± 0.1 and 28.5 ± 0.8 g, have been previously described.^13^

### Sleep/wake recordings

In Study 2, the sleep phenotype of DTA mice with intact, partial, and full ablations of the orexin neuronal field were determined by piezoelectric recording of gross body movement and breath rate (PiezoSleep version 2.11, Signal Solutions, Lexington, KY). By capturing a detailed respiratory measure in addition to locomotor activity, the highly sensitive piezo sensor enables quiet wakefulness to be distinguished from sleep using relevant physiology.^14–17^ The PiezoSleep apparatus consisted of individual polycarbonate cages, each of which contained a polyvinylidene difluoride piezo sensor beneath a thin plastic shield. The sensor covered the entire area of the cage floor and was connected to a single channel tube amplifier (RoMo version 1.2, Signal Solutions, Lexington, KY). Each amplifier fed into a digital acquisition device connected to a computer for data collection and sleep/wake determination as previously described.^14,15,17^ Briefly, signals were amplified and filtered between 0.5 and 10 Hz and analog-to-digital converted at 120 Hz sampling rate. The frequency, amplitude, and peak spectral energy of the signal were evaluated in 2 second increments over a tapered, 8 second window to automatically calculate a sleep/wake decision statistic by simple linear discriminant analysis. Sleep was classified if signals exhibited a periodicity with a consistent, low relative amplitude at a frequency in the typical breathing range (1-4 Hz). Wakefulness was associated with high amplitude signals with variable frequency and broad spectral energy. Signal features related to these properties were extracted and combined to obtain a decision statistic for each interval. Over time, the decision statistics per increment accumulated in a bimodal distribution, and the saddle point of the distribution was determined using an adaptive threshold for the classification of sleep and wake. Bouts of contiguous intervals of sleep or wake were defined to terminate when an interval, equal to the minimum bout length of 30 s, included less than 50% of the targeted state. This method of automated, unsupervised classification of sleep/wake by piezoelectric monitoring in mice has an accuracy of 90% vs. manual EEG scoring^17^ and has been used to characterize the sleep/wake phenotype of numerous transgenic mouse lines, including models of neuropathology with sleep fragmentation^18,19^ and the unconsolidated sleep/wake of narcoleptic Atax mice.^13^

Piezoelectric experiments began by loading mice into individual PiezoSleep cages that contained standard rodent bedding with food and nesting material from each mouse’s home cage. Mice were permitted at least 24 h of acclimation before 42-48 h of experimental data were sampled beginning at ZT6 (14:00). Prior to experimental recording and between recordings on Study Weeks 4 and 7, mice were housed individually in standard cages with nesting material in a different room.

### Experimental design, analysis and statistics

Study 1 employed a between-subjects design to test the interaction of genotype/DOX chow status (WT/standard chow (DOX(−)), DTA mice/DOX(+), or DTA mice/DOX(−) 2d) and sex (male or female) during animal shipment on body weight and orexin-A positive cell counts.

In Study 2, sleep percentages across 24 h and histograms of the percent time spent asleep or awake in bouts from 0.5 to ≥ 64 min were calculated using SleepStats version 2.11 (Signal Solutions, Lexington, KY). Based on the circadian distribution of mean sleep and wake bout durations per h, and on the increased propensity for cataplexy during the early dark period,^1^ sleep and wake data from ZT12-18 were used to differentiate mice based on their ability to maintain long bouts of wakefulness at that time of day. The PiezoSleep narcolepsy screen was comprised of scatterplots of time spent in short sleep bout durations (SBD, < 8 min) vs. long wake bout durations (WBD, ≥ 32 min) during ZT12-18 and was used to evaluate the severity of narcolepsy symptoms in individual mice. The ratio of time spent in long WBD:short SBD represented a “wake-maintenance score” per individual mouse; the lower the ratio, the greater the severity of murine narcolepsy symptoms.^13^ Study 2 used a between-subjects design to investigate the interaction of DOX chow status (DOX(+)7wk, DOX(−)8d (RD)6wk, DOX(+)6wk DOX(−)8d, or DOX(−)7wk) and sex on orexin-A positive cell counts and wake-maintenance scores. A between-within design was used to evaluate the interaction of DOX chow status and repeated measures of circadian time, bout durations bins, and study week recordings.

Statistics were performed using GraphPad Prism version 7.03 (La Jolla, CA). Unless otherwise noted, all data are presented as the mean ± S.E.M. Data were tested for normality using the Shapiro-Wilk test and for equal variances using the Brown-Forsythe test, and if they passed (α = 0.05), parametric inferential statistical tests were applied. If they did not pass these tests, nonparametric statistical tests were used. Grubbs test (α = 0.01) identified an outlier mouse in the DOX(+)7wk group that was subsequently removed from all sleep/wake analyses (time spent in short SBD was 2.6x the value of the upper end of the interquartile range and time spent in long WBD at 0% was > 1.5x beyond the lower end of the interquartile range). In Study 1 body weight changes and orexin-A positive cell counts were compared between groups using two-way analysis of variance (ANOVA) on factors “genotype/DOX chow status” and “sex.” In Study 2, orexin-A positive cell counts in DTA mice were compared between DOX chow groups using the Kruskal-Wallis test because of unequal variance. Sleep time across the day was compared between DTA mice on different DOX chow schedules using two-way mixed-model ANOVA on factors “DOX chow status” (between subjects) and “6-h bin” (within subjects). One-way ANOVA was used to compare the 24 h total percentages of sleep between DOX groups. For the PiezoSleep narcolepsy screen, the relationship between time spent in long WBD and short SBD was determined using Pearson correlation, and the ratio of these variables (long WBD:short SBD, i.e., wake-maintenance score) vs. orexin-A positive cell counts, was determined using Spearman correlation (unequal variance for cell counts). Comparison of the long WBD:short SBD ratio between DOX groups was made using one-way ANOVA. Pearson correlations were used to evaluate the stability of sleep measures (daily sleep amount, time in long WBD, time in short SBD, long WBD:short SBD) between Study Weeks 4 and 7. For each analysis in which significance was indicated (α = 0.05), contrasts between relevant factor levels (genotype/DOX chow status, sex, or time bin, as appropriate) were detected *post hoc* with Bonferroni’s multiple comparisons test (for ANOVA), or multiple comparisons were corrected by controlling the False Discovery Rate using the two-stage step-up method (for Kruskal-Wallis).

## Results

### Histology

In study 1, to determine how 48 h animal shipping conditions might affect DOX chow consumption and therefore orexin neurodegeneration, body weight changes during transit and orexin-A positive cell counts were compared between groups. DTA mice that had been switched to standard chow (DTA DOX(−)2d group) lost 6.7 ± 0.6% (15 males) and 2.4 ± 1.0% (15 females) of their body weight during transit, while DTA mice kept on DOX chow (DOX(+) group) lost 3.4 ± 0.8% (15 males) or gained 0.78 ± 1.0% (15 females) of their body weight. Although males lost a greater percentage of their body weight during transit than females (main effect for “sex,” *F*(1, 56) = 25, *P* < 0.0001) and all DTA mice in the DOX(−)2d group lost more weight than those maintained on DOX(+) (main effect for “DOX status,” *F*(1, 56) = 14.5, *P* = 0.0004), there was no interaction of these factors (*F*(1, 56) = 0.009, *n.s*.). Orexin-A positive cell counts per sex for WT mice and DTA mice in the DOX(+) group and DOX(−)2d group and are shown in Table 1. There was a trend for DTA mice in the DOX(−)2d group to lose 9-13% of their orexin-A positive neurons. While factors “sex” and “genotype/DOX chow status” interacted to determine the number of orexin-A positive neurons present (*F*(2, 18) = 3.83, *P* = 0.04), no contrasts were detected.

**Table 1.** Orexin-A+ cell counts in wild type (WT) and *orexin/tTA;TetOdiphtheria toxin A* (DTA) mice

|  | *N* | WT DOX(-) | DTA DOX(+) | DTA DOX(-) 2d |
| --- | --- | --- | --- | --- |
| Males | 4 | 438 ± 37 | 487 ± 29 | 466 ± 15 |
| Females | 4 | 491 ± 25 | 483 ± 9.6 | 379 ± 28 |
| Both sexes | 8 | 464 ± 23 | 485 ± 14 | 423 ± 22 |

Data are the mean ± S.E.M.
DOX(+), continuous doxycycline chow; DOX(-), standard chow
Two-way analysis of variance:
Interaction: $F(2, 18) = 3.83$ , $P = 0.04$ , no contrasts detected
Main effect, genotype/chow: $F(2, 18) = 3.12$ , $P = 0.07$
Main effect, sex: $F(1, 18) = 0.357$ , *n.s.*

In Study 2, orexin-A positive cell loss was quantified in DTA mice after DOX(−) protocols that were designed to produce partial and full ablation of the orexin field as compared to DTA mice maintained on DOX(+). Differences in the size of partial ablations created during the first week vs. the last week of the study were also assessed. Representative images of the tubural region of the hypothalamus (Figure 2A) and cell counts (Figure 2B) indicate the DOX(−) protocols induced orexin-A positive cell death as expected (*H* = 21.2 (3, *N* = 25), *P* < 0.0001). DTA mice maintained on normal chow for the duration of study (DOX(−)7wk) lost 97.4% (13.5 ± 2.5 cells remained) of their orexin-A positive neurons compared to DTA mice maintained on DOX(+) for 7 weeks (525 ± 17 cells). Partial ablation of the orexin neuronal field that was initiated during Study Week 1 (DOX(−)8d (RD)6wk group) created more cell loss (71.0%, 153 ± 21 cells remained) than the same duration of DOX(−) that occurred during Study Week 7 (DOX(+)6wk DOX(−)8d group, 56.6% cell loss, 228 ± 15 cells remained). The distribution of orexin-A positive cells in males vs. females in each DOX chow group is shown in Supplemental Figure 1.

**Figure 2.**
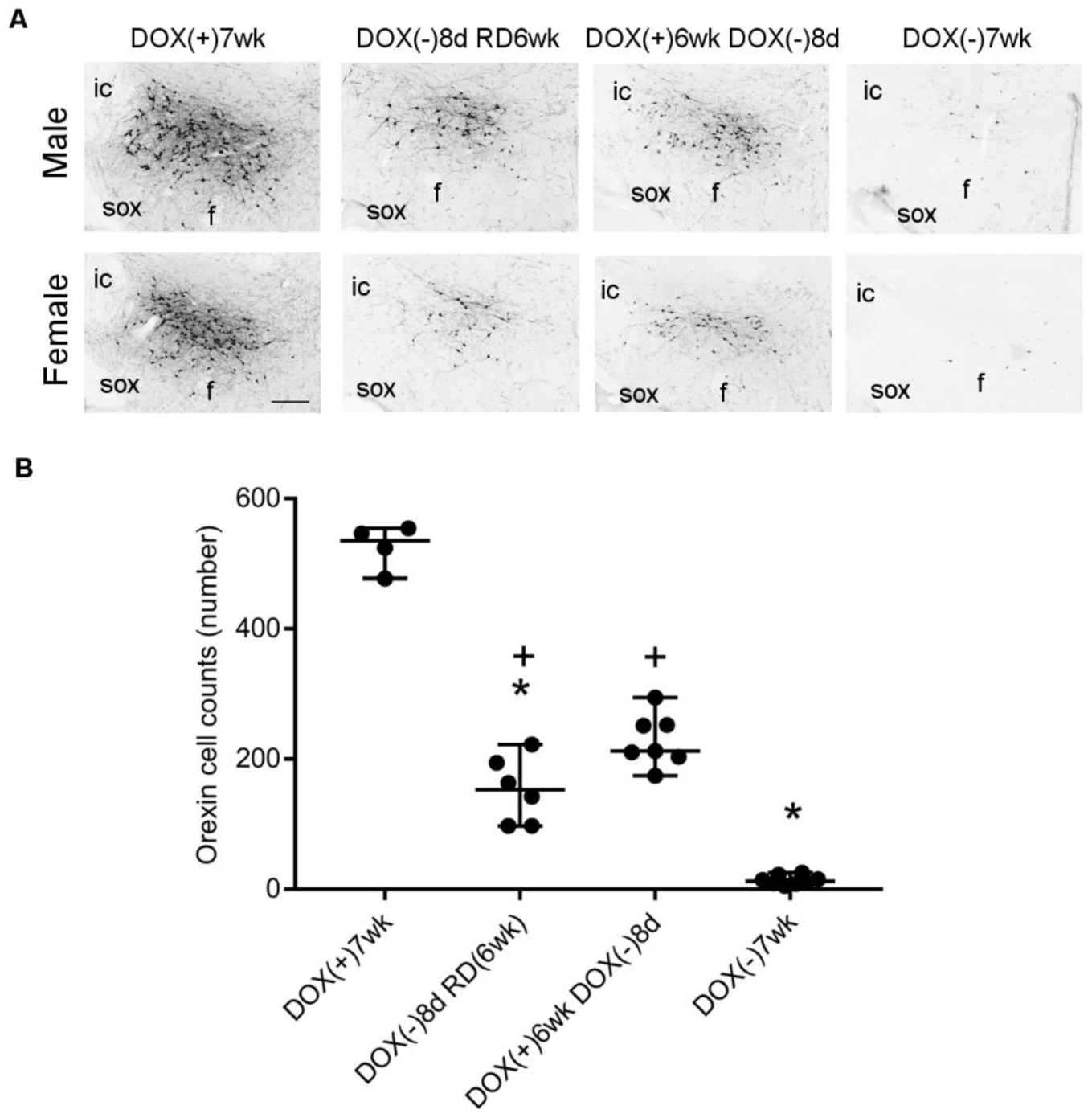
Loss of orexin-A positive neurons in *orexin/tTA; TetO diphtheria toxin A* (DTA) mice in Study 2. Representative images of Orexin-A+ cells in the posterior lateral hypothalamus **(A)** of DTA colony mates maintained on doxycycline for 7 weeks (DOX(+)7wk, *N* = 2 males, 2 females), with removal of doxycycline for 8 days and resumption of DOX(+) for 6 wk (DOX(−)8d (RD)6wk, *N* = 4 males, 2 females), with removal of doxycycline 8 days prior to sacrifice (DOX(+)6wk DOX(−)8d, *N* = 4 males, 3 females), or after 7 weeks without doxycycline (DOX(−)7wk, *N* = 4 males, 4 females). Images taken at 10x magnification (bar = 250 μm), approximately -1.5 mm from Bregma; Fornix (f), internal capsule (ic), supraoptic decussation (sox). Orexin-A+ cell counts from individual mice in each group **(B)**, delineated with the median and 95% confidence intervals. Kruskal-Wallis with multiple comparisons correction: * P < 0.05 vs. DOX(+)7wk, ^+^P < 0.05 vs. DOX(−) 7wk.

### Sleep/wake phenotypes

The relationship between partial orexin neurodegeneration and the quantity and architecture of sleep/wake was evaluated in DTA mice after ablation was permitted by DOX(−)8d (RD)6wk (71.0% orexin-A positive cell loss) or by DOX(+)6wk DOX(−)8d (56.6% orexin-A positive cell loss) vs. DTA mice maintained as orexin-intact on DOX(+) or with 97.4% orexin-A positive cell loss from DOX(−) for 7 weeks. The percentage of time spent asleep over 24 h or within 6-h bins across the day did not depend on the extent of orexin ablation (Figure 3), *F*(3, 20) = 0.24, *n.s*. and *F*(9, 57) = 1.18, *n.s*., respectively.

**Figure 3.**
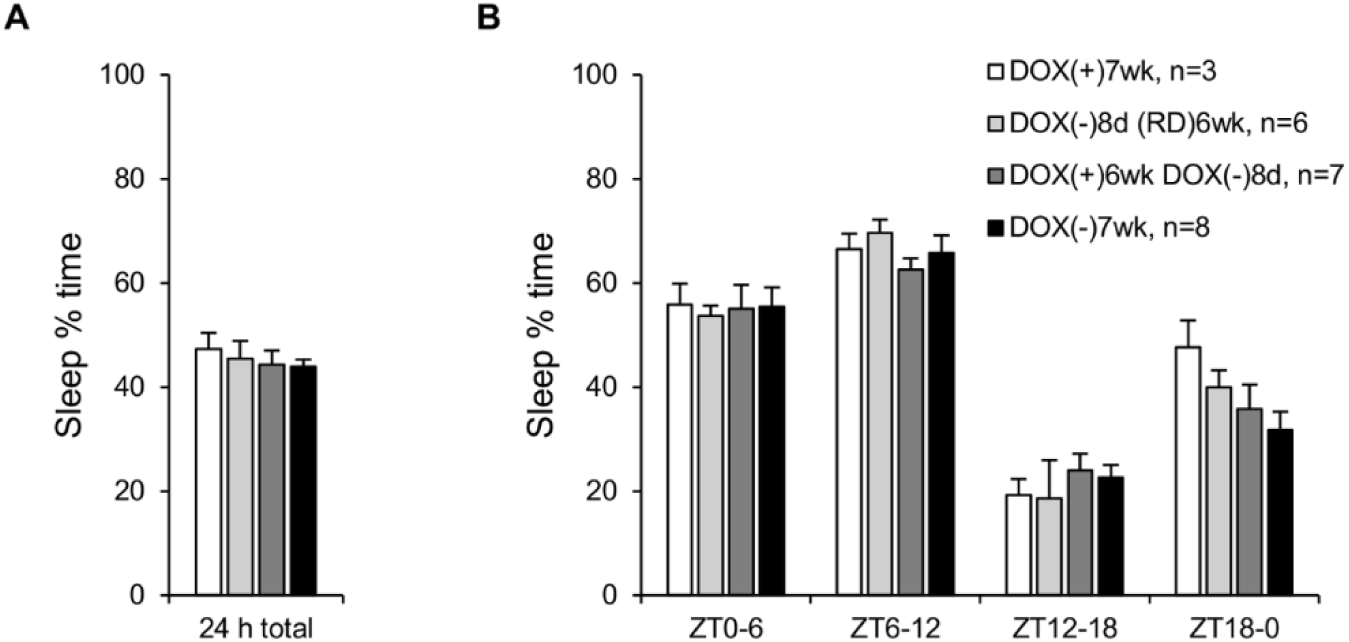
Orexin neurodegeneration does not affect the daily amount nor distribution of sleep time in *orexin/tTA; TetO diphtheria toxin A* (DTA) mice. Percent of time spent asleep (mean ± S.E.M.) per 24 h **(A)** or across the day in 6-h time bins **(B)**. DTA colony mates were maintained on doxycycline for 7 weeks (DOX(+)7wk, *N* = 2 males, 1 female) **(white)**, with removal of doxycycline for 8 days and resumption of DOX(+) for 6 wk (DOX(−)8d (RD)6wk, *N* = 4 males, 2 females) **(light gray)**, with removal of doxycycline 8 days prior to sacrifice (DOX(+)6wk DOX(−)8d, *N* = 4 males, 3 females) **(dark gray)**, or after 7 weeks without doxycycline (DOX(−)7wk, *N* = 4 males, 4 females) **(black)**.

The severity of murine narcoleptic symptoms in DTA mice with full orexin-A positive cell loss (DOX(−)7wk) vs. those with partial ablation of the orexin field (DOX(−)8d (RD)6wk and DOX(+)6wk DOX(−)7wk) or those with intact orexin neurons (DOX(+)7wk) were compared using the PiezoSleep narcolepsy screen (Figure 4). As expected, the negative correlation between time spent in short SBD to long WBD was significant, and mice with full orexin-A positive cell loss tended to cluster in a manner reflective of their low ratio of time in long WBD:short SBD (Figure 4A). The wake-maintenance scores (ratio of time spent in long WBD:short SBD) correlated with orexin-A positive cell counts and further distinguished DTA mice on the basis of their narcoleptic phenotype (Figure 4B). The DTA mice with 97.4% orexin-A positive cell loss resembled Atax mice (96.6 % cell loss, Figure 4B) and exhibited a lower wake-maintenance score than DTA mice maintained on DOX(+) with intact orexin neurons (Figure 4C), *F*(3, 20) = 14.46, *P* < 0.0001. Mice with partial ablation of the orexin field clustered intermediary to DOX(+)7wk and DOX(−)7wk mice. The wake-maintenance score was 3.8x greater in mice that received the partial ablation during Study Week 1 (DOX(−)8d (RD)6wk) than in mice with near complete orexin-A positive cell loss (DOX(−)7wk) and was not significantly different from the ratio observed in orexin-intact mice. The wake-maintenance score in mice that received partial ablation in the week before sacrifice (DOX(+)6wk DOX(−)8d) was lower than in orexin-intact mice and was not significantly different from the ratio observed in DOX(−)7wk mice. Wake-maintenance scores in individual male vs. female DTA littermates by orexin-A positive cell counts are depicted in Supplemental Figure 2.

**Figure 4.**
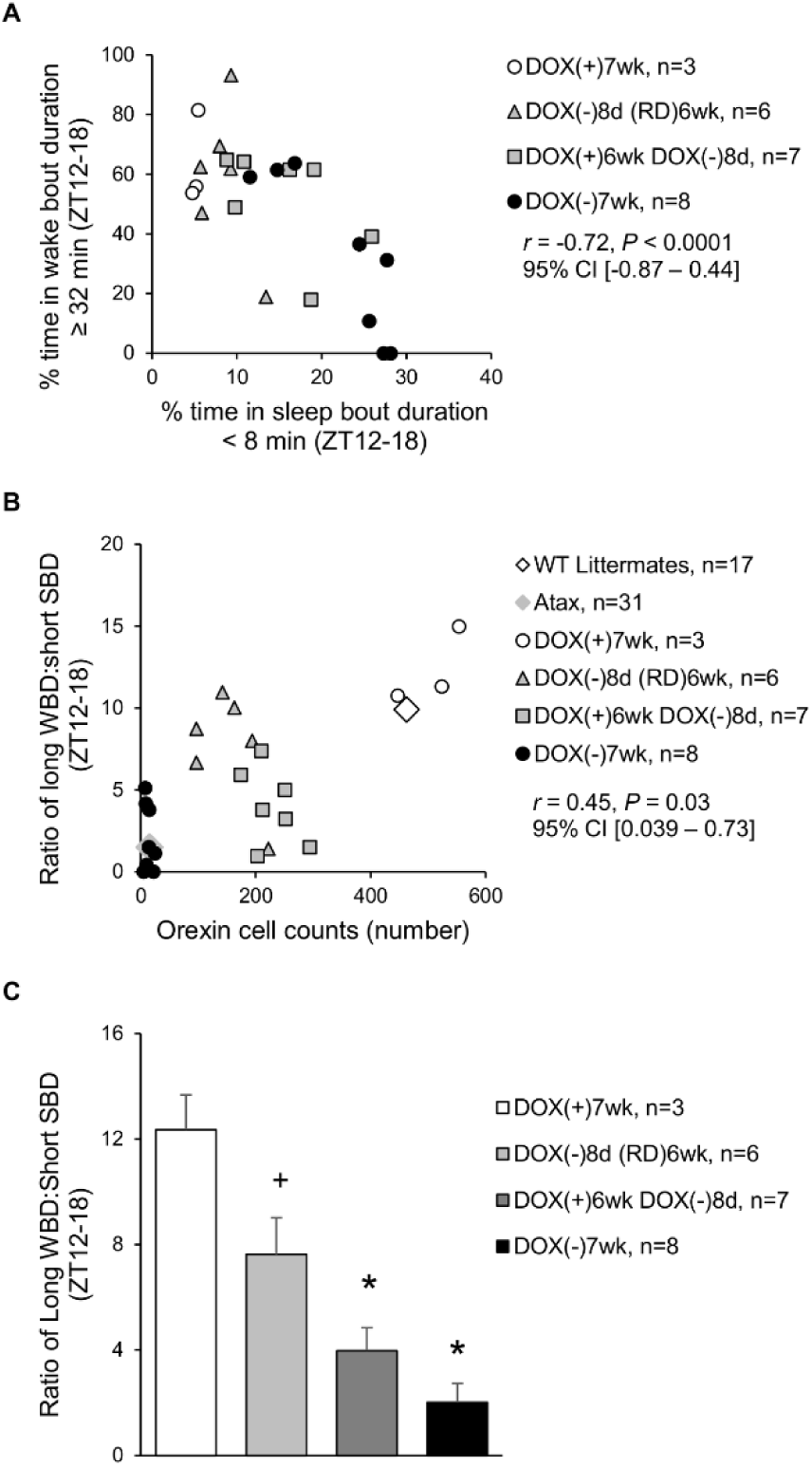
Partial ablation of the orexin field induces a sub narcoleptic phenotype as revealed in the PiezoSleep narcolepsy screen. Percent of time spent in short sleep bouts (< 8 min) vs. long wake bouts (≥ 32 min) from ZT12-18 per individual *orexin/tTA; TetO diphtheria toxin A* (DTA) mouse **(A)**. Ratio of time in long wake bouts to short sleep bouts from ZT12-18 vs. orexin-A+ cell count per individual **(B)** and mean (± S.E.M.) per doxycycline/orexin ablation group **(C)**. Male and female DTA colony mates were maintained on doxycycline for 7 weeks (DOX(+)7wk, *N* = 4 males, 1 female) **(white circles and bar)**, with removal of doxycycline for 8 days and resumption of DOX(+) for 6 wk (DOX(−)8d (RD)6wk, *N* = 4 males, 2 females) **(gray triangles, light gray bar)**, with removal of doxycycline 8 days prior to sacrifice (DOX(+)6wk DOX(−)8d, *N* = 4 males, 3 females) **(gray squares, dark gray bar)**, or after 7 weeks without doxycycline (DOX(−)7wk, *N* = 4 males, 4 females) **(black circles and bar)**. Male *orexin/ataxin-3* (Atax) mice **(gray diamond)** and male wild type colony mates **(white diamond)** are plotted as group means (paired sleep and cell count data from individuals not available) for reference and were not included in the statistical analysis. Data from DTA mice were evaluated by Pearson correlation **(A)**, Spearman correlation (because of unequal variance for cell counts) **(B)**, and one-way analysis of variance with Bonferroni’s multiple comparisons **(C)**, *P* < 0.05: * vs. DOX(+)7wk, ^+^vs. DOX(−)7wk. Supplemental data **(Supplemental Figure 2)** show ratio of time in long wake bouts to short sleep bouts from ZT12-18 vs. orexin-A+ cell count per individual by sex.

The stability of the sleep/wake phenotype over time in DTA mice, in which ablation of the orexin neuronal field began in Study Week 1, and in orexin-intact controls, was evaluated by correlative comparison of data collected during Study Weeks 4 and 7. As expected, the amount of sleep per 24 h strongly correlated between study weeks (Figure 5A). Measures of sleep/wake fragmentation during ZT12-18 from the narcolepsy phenotyping screen showed more variation between Study Weeks 4 and 7 than the 24-h total sleep times. The percentage of time spent in WBD ≥ 32 min (Figure 5B), in SBD < 8 min (Figure 5C), and the ratio of these variables (Figure 5D) also strongly correlated between Study Weeks 4 and 7.

**Figure 5.**
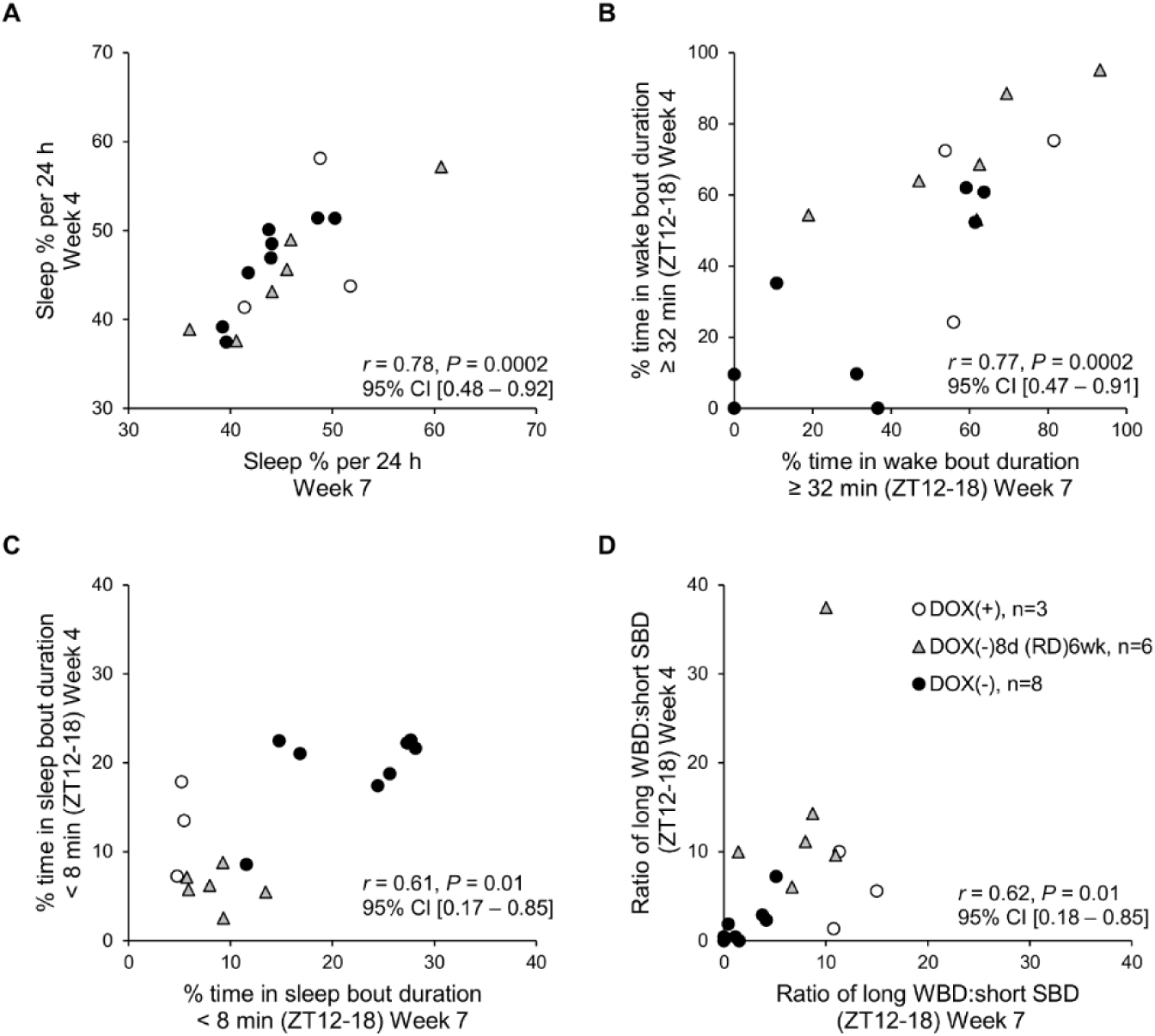
Stability of sleep phenotype in *orexin/tTA; TetO diphtheria toxin A* (DTA) mice over time. Correlations of data sampled on Study Week 4 vs. 7 for the percent of time spent: asleep per 24 h **(A)**, in long wake bout durations (WBD, ≥ 32 min) **(B)**, or short sleep bout durations (SBD, < 8 min) **(C)**, or for the ratio of time spent in long WBD:short SBD from ZT12-18 **(D)**. DTA colony mates were maintained on doxycycline for 7 weeks (DOX(+)7wk, *N* = 2 males, 1 female) **(white circles)**, with removal of doxycycline for 8 days and resumption of DOX(+) for 6 wk (DOX(−)8d (RD)6wk, *N* = 4 males, 2 females) **(gray triangles)**, or after 7 weeks without doxycycline (DOX(−)7wk, *N* = 4 males, 4 females) **(black circles)**. The group in which DOX(−)8d occurred during study week 7 were not included in the analysis because of the change in DOX status between sleep sampling time points. Data were evaluated by Pearson correlation **(A, B)** or by Spearman correlation, because data from Study Week 4 did not pass the normality test **(C, D)**.

## Discussion

For the first time, sleep/wake phenotypes and orexin-A positive cell counts have been determined in the same individual DTA mice after partial ablation of the orexin neuronal field and sleep/wake measurement by piezoelectric monitoring. First, the effect of animal shipment conditions, which may disrupt the normal consumption of DOX chow, on the integrity of the Tet Off system was assessed. The DTA mice in transit for 48 h and maintained on DOX chow (200 μg/g chow) did not lose orexin-A positive neurons despite weight loss, and presumably reduced DOX chow consumption. Because DOX(−)2d during transit only induced 9-13% orexin-A positive cell loss, in Study 2, DTA mice were subjected to DOX(−) 8d to produce more substantial neurodegeneration. Partial ablations of the orexin field that were created during the first week of study (DOX(−)8d (RD)6wk group) were 14% larger than the partial ablations induced by the same duration of DOX(−) that occurred in the last week of study (DOX(+)6wk DOX(−)8d group), suggesting orexin cell death continued despite the resumption of DOX chow for 6 weeks. Despite these larger partial ablations (71.0% orexin-A positive cell loss), the wake-maintenance scores of mice in the DOX(−)8d (RD)6wk group resembled those of orexin-intact mice more than those of mice with near complete (≥ 97%) orexin-A positive neurodegeneration. Conversely, mice with smaller partial ablations (56.6% orexin-A positive cell loss) in the DOX(+)6wk DOX(−)8d group had lower wake-maintenance scores that resembled those of mice with near complete orexin-A positive cell loss.

Collectively, these results suggest that compensatory wakefulness, perhaps through the histaminergic system,^20,21^ develops over time in mice that have some critical portion of their orexinergic system remaining after partial ablation. These compensatory changes may occur mostly within the first three weeks, because sleep/wake phenotypes remained stable within individual mice between Study Weeks 4 and 7.

An important factor relevant to the interpretation of the neurodegeneration results in the current study vs. previous work concerns the different concentrations of DOX chow employed. In Study 1, DOX(−)2d resulted in ~5x less orexin-A positive cell loss than anticipated based on the rate of neurodegeneration previously reported.^12^ Doxycycline in the current study was provided at twice the concentration as was used previously for an estimated daily dose of 24 mg/kg. An approximately 10-fold lower daily dose of DOX, as supplied by DOX (10 μg/mL) in drinking water, has been recommended as the optimal concentration for manipulation of gene expression within 1-2 days using the TetO system due to the very slow kinetics of washout and reactivation of the transgene.^22^ Although the pharmacokinetics of DOX used in the current study is unknown, the dose used was ~16-fold lower than the dose achieved with DOX (2 mg/mL), which has been shown to cause residual suppression of gene expression after 8 weeks washout.^23^ In the current study, the smaller than expected ablation sizes and the continued neurodegeneration that was observed for weeks past the resumption of DOX may be related to the slow clearance of the DOX (200 μg/g) diet.

Orexin insufficiency to varying degrees, as measured through either CSF orexin-A levels or postmortem histology, has been associated with a variety of neurological conditions in addition to narcolepsy and expands the clinical relevance of the DTA mouse. Orexin cell loss of 27% in Huntington’s disease,^24^ 23-62% in Parkinson’s disease,^25,26^ and 69% in multiple system atrophy^27^ has been demonstrated despite normal levels of CSF orexin-A in these patient populations.^28–31^ This discrepancy suggests that the residual orexin neurons compensate for reduced orexin connectivity with increased peptide release. Compensation has been observed in rats with hypothalamic orexin-B-saporin lesions; 73% orexin cell loss resulted in only 50% loss of CSF orexin-A.^32^ If this relationship can be extrapolated to DTA mice, then it can be speculated in the current study that mice with partial ablations may have had CSF orexin-A levels that were 40-50% less than those in orexin-intact controls. DTA mice with ≥ 97% orexin neurodegeneration may have had ≥ 66% loss of CSF orexin-A. This level of remaining CSF orexin-A (~1/3 of orexin-intact controls) is very close to the cutoff value (30% of controls or 110 pg/mL) that is most predictive of narcolepsy with cataplexy in the human diagnostic test.^7^ Future research to test these relationships between orexin cell loss and CSF orexin-A levels, and thereby increase the translational value of the partially ablated DTA mouse, will require the development of a highly sensitive assay to detect orexin-A levels in small volume (10 μL) CSF samples from individual mice, as current assays require pooling samples to obtain 5x that volume.^9^

Narcolepsy type 2 is a heterogeneous disease entity with uncertain pathophysiology that makes clinical diagnosis and animal modeling challenging. Diagnosis of Na-2 requires: EDS, a positive score on the MSLT (sleep latency ≤ 8 min, and ≥ 2 sleep onset REM periods), an absence of cataplexy, and differentiation from idiopathic hypersomnia by signs of REM sleep boundary dysregulation.^33^ The majority of patients with Na-2 have normal CSF orexin-A levels,^7^ but 8% of patients with narcolepsy without cataplexy were found to have intermediate CSF orexin-A levels that were above the Na-1 cutoff value and below the normal level of ≥ 200 pg/mL.^34^ Seven out of 10 patients with intermediate CSF orexin-A levels were found to have atypical or no cataplexy.^6^ In patients with intermediary orexin-A levels, 18% developed cataplexy years after the original narcolepsy-without-cataplexy diagnosis.^34^ These data, together with the intermediary number of orexin cells found postmortem from one patient with narcolepsy without cataplexy,^5^ support the hypothesis that at least some cases of Na-2 lie on a continuum between Na-1 and idiopathic hypersomnia. The current study offers the first steps toward development of a mouse model of Na-2 based on this hypothesis that partial disruption of orexinergic signaling results in pathological sleepiness associated with less severe narcolepsy. However, the current study is limited by the lack of cataplexy and REM sleep measurements. Cataplexy assessment would be necessary to fully differentiate whether the phenotypes observed in DTA mice with partial ablations of the orexin field translate as Na-1 or Na-2. REM sleep timing may further distinguish subtle differences between DTA mice with different sizes of partial ablations. Measuring cataplexy requires invasive, time-consuming, and expensive procedures, and as such, would be the next step after establishment of protocols for creating specific ablation sizes that are associated with the mouse equivalent of excessive daytime sleepiness.

In conclusion, a sub narcoleptic phenotype was modeled in DTA mice to the extent that a range of wake-maintenance scores was found to be associated with different sizes and durations of orexin-A positive cell loss. The relationship between partial orexin neurodegeneration and sleep/wake phenotype depends on the dynamics of progressive orexin cell loss and the development of a compensatory wake response. Further work is recommended to titrate the DOX(−) RD protocol for the induction of stable 80-90% orexin neurodegeneration. This amount of orexin cell loss is predicted to distinguish partially ablated DTA mice from orexin-intact controls without permitting cataplexy. The establishment of a mouse model based on partial ablation of orexin neurons will be critical for efficacy tests of orexin modulator therapies for narcolepsy and other diseases characterized by orexin insufficiency.

## Acknowledgements

This work was funded by Reset Therapeutics and supported in part by the Michael J. Fox Foundation (grant number 11432 to Dr. Humphries). The content is solely the responsibility of the authors and does not necessarily represent the official views of the Michael J. Fox Foundation. We thank Dr. Thomas Scammell (Harvard University) and Dr. Takeshi Sakurai (University of Tsukuba) for the donation of the orexin/ataxin-3 mice to Jackson Labs, the staff at Neuroscience Associates for the histology, Tate York (NeuroScience Associates) for the image analysis, Dr. Kevin Donohue and Dr. Bruce O’Hara (Signal Solutions and University of Kentucky) for assistance with PiezoSleep, and Tod Steinfeld and Dr. Bruce Clapham (Reset Therapeutics) for helpful discussions.

## Abbreviations

ANOVA: analysis of variance
Atax: *orexin/ataxin-3*
CSF: cerebrospinal fluid
DOX: doxycycline
DOX(+)6wk DOX(−)8d: DOX chow for 6 weeks followed by standard chow for 8 days
DOX(+)7wk: DOX chow for 7 weeks
DOX(−)7wk: standard chow for 7 weeks
DOX(−)8d (RD)6wk: standard chow for 8 days followed by resumed DOX chow for 6 weeks
*DTA*: *orexin-tTA; TetO diphtheria toxin A*
EDS: excessive daytime sleepiness
EEG: electroencephalography
EMG: electromyography
f: fornix
ic: internal capsule
LD12:12: 12 h light period, 12 h dark period
MSLT: multiple sleep latency test
MWT: maintenance of wakefulness test
Na-1: narcolepsy type 1
Na-2: narcolepsy type 2
NREM: non-rapid-eye-movement
PBS: phosphate-buffered saline
REM: rapid-eye-movement
RM ANOVA: repeated-measures analysis of variance
SBD: sleep bout duration
S.E.M.: standard error of the mean
Sox: supraoptic decussation
TetO: Tet Off
tTA: tetracycline transactivator
WBD: wake bout duration
WT: wild type
ZT: zeitgeber time

## Supplemental Data

**Supplemental Figure 1.**
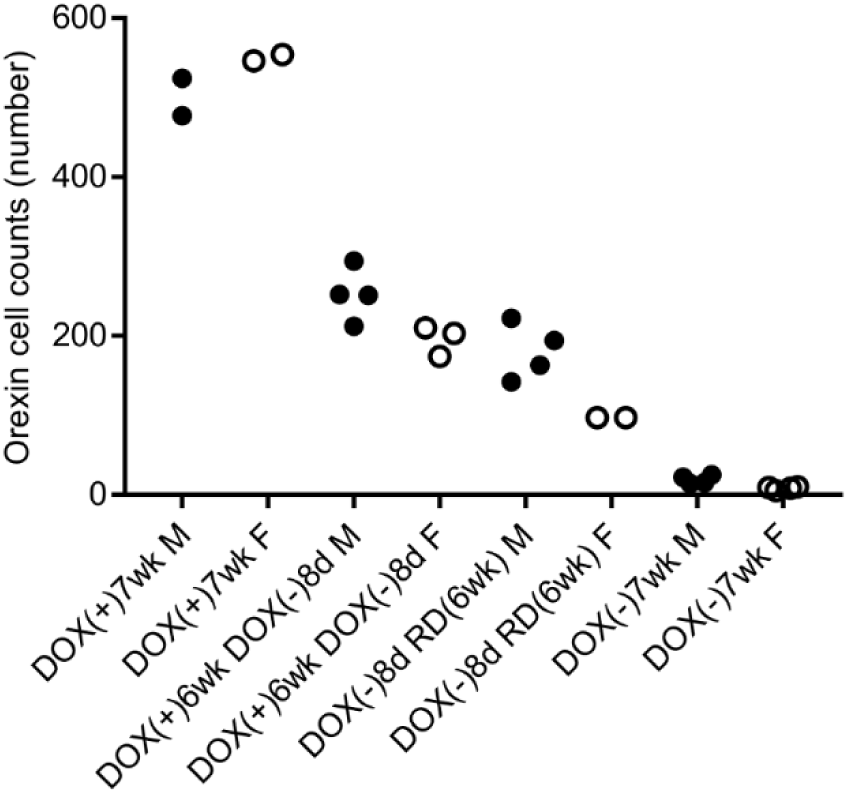
Orexin-A+ cell counts in individual male (M) vs. female (F) *orexin/tTA; TetO diphtheria toxin A* (DTA) mice in Study 2. DTA colony mates were maintained: on doxycycline for 7 weeks (DOX(+)7wk), with removal of doxycycline 8 days prior to sacrifice (DOX(+)6wk DOX(−)8d), with removal of doxycycline for 8 days and resumption of DOX(+) for 6 wk (DOX(−)8d (RD)6wk), or after 7 weeks without doxycycline (DOX(−)7wk).

**Supplemental Figure 2.**
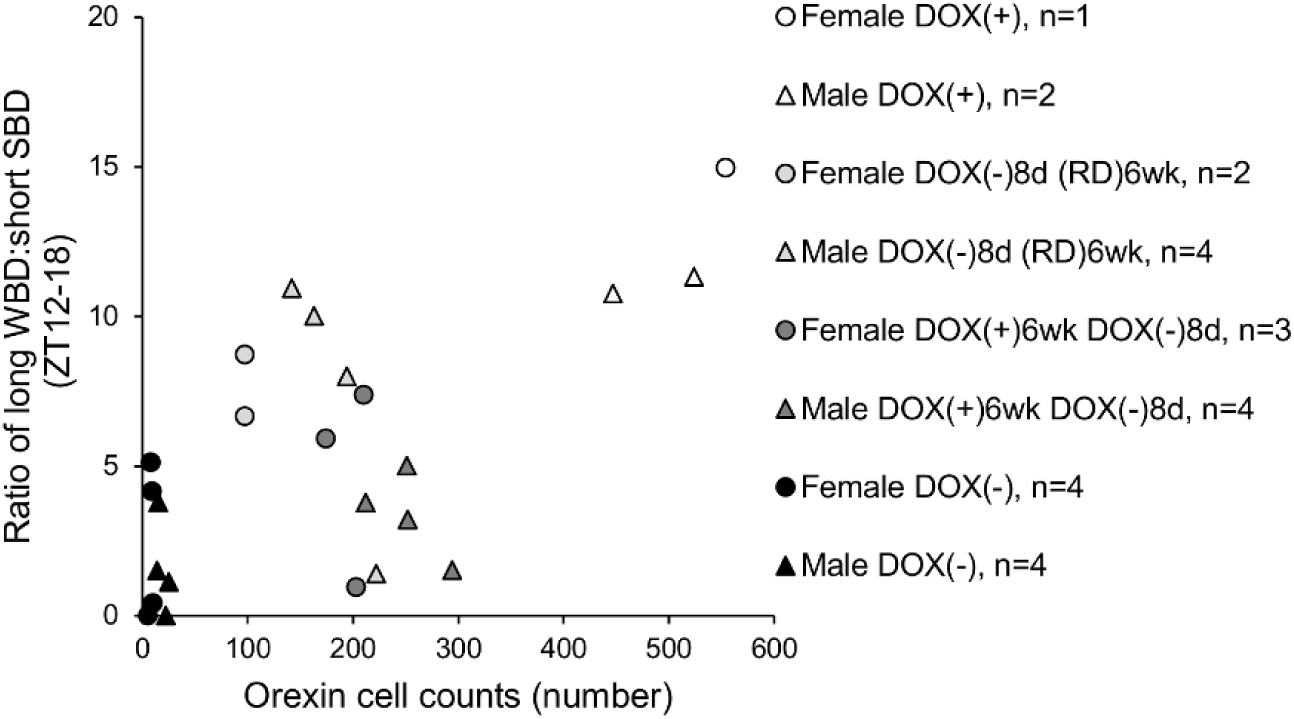
Narcoleptic phenotypes in male vs. female *orexin/tTA; TetO diphtheria toxin A* (DTA) mice after partial, full, or no ablation of the orexin neuronal field. Ratio of time spent in long wake bout durations (WBD) to short sleep bout durations (SBD) from ZT12-18 vs. orexin-A+ cell count per individual (expressed as percent of orexin-intact control mean per sex). Male **(triangles)** and female **(circles)** DTA colony mates were maintained on doxycycline for 7 weeks (DOX(+)7wk) **(white symbols)**, with removal of doxycycline for 8 days and resumption of DOX(+) for 6 wk (DOX(−)8d (RD)6wk) **(light gray symbols)**, with removal of doxycycline 8 days prior to sacrifice (DOX(+)6wk DOX(−)8d) **(dark gray symbols)**, or after 7 weeks without doxycycline (DOX(−)7wk) **(black symbols)**.

